# Increased Ethanol Intake is Associated with Social Anxiety in Offspring Exposed to Ethanol on Gestational Day 12

**DOI:** 10.1101/2020.03.13.991562

**Authors:** Marvin R. Diaz, Julia M. Johnson, Elena I. Varlinskaya

**Affiliations:** Department of Psychology, Center for Development and Behavioral Neuroscience Binghamton University, Binghamton NY13902, United States; Developmental Exposure Alcohol Research Center, Baltimore MD21201; Binghamton NY 13902; Syracuse NY13210, United States

**Keywords:** Prenatal, fetal, alcohol, anxiety, social, ethanol intake

## Abstract

Prenatal alcohol exposure (PAE) can result in physical, cognitive, and neurological deficits termed Fetal Alcohol Spectrum Disorder (FASD). Deficits in social functioning associated with PAE are frequently observed and persist throughout the lifespan. Social impairments, such as social anxiety, are associated with increased alcohol abuse, which is also highly pervasive following PAE. Yet, the relationship between PAE-induced social alterations and alcohol intake later in life is not well understood. In order to test this relationship, we exposed pregnant female Sprague Dawley rats to a single instance of PAE on gestational day 12 and tested offspring in adulthood (postnatal day 63) in a modified social interaction test followed by alternating alone and social ethanol intake sessions. Consistent with our previous findings, we found that, in general, PAE reduced social preference (measure of social anxiety-like behavior) in female but not male adults. However, ethanol intake was significantly higher in the PAE group regardless of sex. When dividing subjects according to level of social anxiety-like behavior (low, medium, or high), PAE males (under both drinking contexts) and control females (under the social drinking context) with a high social anxiety phenotype showed the highest level of ethanol intake. Taken together, these data indicate that PAE differentially affects the interactions between social anxiety, ethanol intake, and drinking context in males and females. These findings extend our understanding of the complexity and persistence of PAE’s sex-dependent effects into adulthood.

## Introduction

The incidence of alcohol drinking and alcohol abuse during pregnancy is staggeringly high, with estimated rates of 10-20% in the US [1] and as high as 50% in Italy [2]. The consequences of prenatal alcohol exposure (PAE) are collectively termed Fetal Alcohol Spectrum Disorders (FASDs) that affect ∼5% of the US population [3], a likely underestimate due to the stigma of disclosing drinking habits during pregnancy [4]. The FASD diagnosis can range from severe mental retardation to more subtle behavioral deficits, or Alcohol Related Neurodevelopmental Disabilities (ARNDs). Some of the most disabling effects of PAE are the high rates of anxiety disorders, estimated at ∼21%, and the numerous impairments in social behavior observed throughout development and into adulthood [5]. Additionally, and equally important, PAE leads to a higher risk for developing substance abuse problems [6-9], with levels as high as 29% in adolescents (12-20.9 years) and 46% in adults (21+ years) with FASD having alcohol or drug abuse problems [reviewed in [6]]. Importantly, alcohol abuse is known to be highly associated with anxiety [10], as a result of both the anxiolytic and social facilitating properties of alcohol [11-13] and neuroadaptations in key overlapping neural circuits [14].

Consistent with neurobehavioral deficits associated with FASD, animal models have recapitulated many of the long-lasting effects of PAE. For example, many groups have shown that PAE across various gestational periods can alter non-social anxiety-like behaviors at various ages [15-27]. In rodents, PAE also leads to increased ethanol intake in offspring across various ages [28-35]. Alterations in social behavior have also been observed in rodents exposed to ethanol *in utero* [36-43]. Importantly, we have shown that a single exposure to a high dose of ethanol on gestational day (G) 12 increases social anxiety-like behaviors in an age- and sex-dependent manner in two strains of rats, Long Evans [44] and Sprague Dawley [27]. Additionally, G12 exposed adolescent offspring exhibited reduced sensitivity to the socially anxiogenic effects of an acute ethanol challenge, and demonstrated social facilitation at low-doses of ethanol known to not alter social behavior in control rats [45].

Despite the numerous studies that have consistently identified behavioral alterations following PAE, and the known associations between anxiety, social behavior, and ethanol use [11-13, 46], no study has assessed both ethanol intake and social behavior within the same subject following PAE. Given the clear link between anxiety and ethanol preference following PAE, the objective of this study was to determine whether G12 PAE would alter ethanol intake in adult offspring in either social- or alone-drinking circumstances and examine the association between social anxiety-like behavior and ethanol intake.

## Methods

### Subjects

Experimental subjects were the offspring of Sprague-Dawley rats that were produced by time mating in our colony at Binghamton University; colony originated from animals purchased through Taconic Inc. (Blooming Grove, PA). All animals were housed in a temperature-controlled (22°C) vivarium and maintained on a 12:12 h light:dark cycle (lights on at 0700 h) and had *ad libitum* access to food (Purina rat chow) and water. For breeding procedures, three adult females were housed with one male in a large plastic cage during a 4-day period. Vaginal smears were collected every day, with the first day of detectable sperm designated as G1 [27, 44, 45].

### Prenatal Alcohol Exposure

As previously described [27, 45], females were injected intraperitoneally (i.p.) with 2.5 g/kg ethanol (20% v/v ethanol in physiological saline) on G12. Two hours later, pregnant females were given a second i.p. injection of 1.25 g/kg ethanol. Control females received two i.p. injections of equivalent volumes of saline. Pregnant dams were undisturbed until parturition (postnatal day (P) 0). Non-injected dams produced offspring later used as social partners. Litters were culled to 10 pups by P2, maintaining a 1:1 sex ratio when possible. Pups remained with their dams in standard plastic home-cages with pine shavings as bedding material. Offspring were weaned on P21 and placed into standard plastic cages with same-sex littermates (2 animals per cage). Maintenance and treatment of animals was in accordance with standard guidelines for animal care established by the National Institutes of Health and approved by the Binghamton University Institutional Animal Care and Use Committee.

### Social interaction test

A modified social interaction test was conducted in young adult (P63) offspring as previously described [27, 45]. All testing was performed under dim light in Plexiglas test apparatuses (45 x 30 x 20 cm; Binghamton Plate Glass, Binghamton, NY) containing clean shavings. Test apparatuses were divided into two equally sized compartments by a clear Plexiglas partition with an aperture (7 x 5 cm) to allow animals to move between compartments. Given the size of the aperture, only one animal was able to move through the aperture at a time. On test day (P63), animals were taken from their home cage and individually placed in the testing apparatus for 30 min. A same age and sex social partner was then introduced for a 10 min testing period. Partners were not socially deprived prior to the test and were experimentally naïve but were unfamiliar with both the apparatus and the experimental animal. Weight differences between test subjects and their partners did not exceed 20 g for adult males and 10 g for adult females, with test subjects always being heavier than their partners. To distinguish experimental animals and social partners during the test, experimental animals were marked with a vertical black line on the back. The behavior of the animals was video recorded during the 10 min test session. All testing procedures were conducted between 0900 and 1100 h under dim light (15-20 lux). The number of crossovers demonstrated by the experimental subject towards, as well as away from, the social partner was measured separately, and a coefficient of social preference/avoidance was then calculated [coefficient (%) = (crossovers to the partner − crossovers away from the partner)/(total number of crosses both to and away from the partner) × 100]. The coefficient of social preference/avoidance is a reliable measure of social anxiety-like alterations, with lower positive and negative values associated with social anxiety-like alterations [47-50]. High positive values of the coefficient are characteristic of low anxiety evident under social test circumstances.

### Ethanol intake

One week after assessment of social interaction, rats were tested for six 30-minute drinking sessions every Monday, Wednesday, and Friday (P70-81) as previously done [51]. Briefly, on alternating drinking days, animals were given access to 10% ethanol in “supersac” (3% sucrose + 0.125% saccharin solution) either alone or in a group of two littermates, with the order of social/alone drinking sessions counterbalanced within each prenatal exposure/sex group. Drinking sessions were conducted in a novel cage, and experimental subjects were not food or water deprived. Sessions were videotaped for later scoring. To determine ethanol intake, bottles containing ethanol + supersac solution were weighed before and after drinking sessions, and time spent drinking for each individual animal (for social drinking) was calculated from videos. After 30 mins, animals were returned to their home cages with their cage mate. Individual intake was calculated by determining proportional time spent drinking for each animal as follows: ml (or g of ethanol) consumed by a pair/total time spent drinking x time spent drinking for an individual animal. Trunk blood samples were immediately taken after the last drinking session to determine blood ethanol concentrations (BECs). Samples were collected in heparinized tubes, rapidly frozen, and maintained at −80°C until analysis of BECs. BECs were quantified using headspace gas chromatography as done previously [45].

### Statistics

Given our previous data that showed sex-dependent effects of ethanol exposure on G12 [27, 44, 45], all analyses were done separately in males and females. Impact of prenatal ethanol exposure on ethanol intake in adulthood was assessed using a 2 (prenatal exposure: Sal, EtOH) x 6 (test day) analysis of variance (ANOVA), with Day treated as a repeated measure. Prenatal exposure-related differences in ethanol intake under social or alone circumstances averaged across the three drinking sessions were analyzed using a 2 (prenatal exposure) X 2 (drinking context) ANOVA, with context treated as a repeated measure. BECs assessed on test day 6 were also analyzed using a 2 (prenatal exposure) X 2 (drinking context) ANOVA.

A tertile split of the coefficient of social preference/avoidance was used within each sex/prenatal condition for determination of animals with low (n=7 per group), medium (n=6/group), and high (n=7/group) social anxiety-like behavior. The influence of social anxiety-like behavior indexed via low social preference on subsequent ethanol intake under social or alone circumstances was analyzed for each prenatal exposure/sex group using separate 3 (level of social anxiety-like behavior: low, medium, high) X 2 (drinking context) mix-factor ANOVAs, with drinking context treated as a repeated measure. In order to avoid inflating the possibility of type II errors on tests with at least 3 factors [52], Fisher’s planned pair-wise comparison test was used to explore significant effects and interactions.

## Results

In males, ethanol intake differed as a function of prenatal exposure [main effect of Exposure, F(1,190) = 7.385, p < 0.01], with males prenatally exposed to ethanol on G12 ingesting more ethanol than their saline-exposed counterparts (see Figure 1, left). Males in both prenatal exposure conditions gradually increased their intake from day 1 to day 4 [main effect of Day, F(5,190) = 29.136, p < 0.0001]. In females, the ANOVA revealed a significant Prenatal Exposure x Test Day interaction, F(5, 190) = 2.290, p < 0.05. Females from both prenatal conditions increased their ethanol intake from test day 1 to test day 3, with no differences evident between ethanol- and saline-exposed animals. The differences between saline- and ethanol-exposed females became apparent on test days 4, 5, and 6, with young adult females exposed to ethanol on G12 demonstrating substantially higher intake than their saline-exposed counterparts (Figure 1, right).

**Figure 1:**
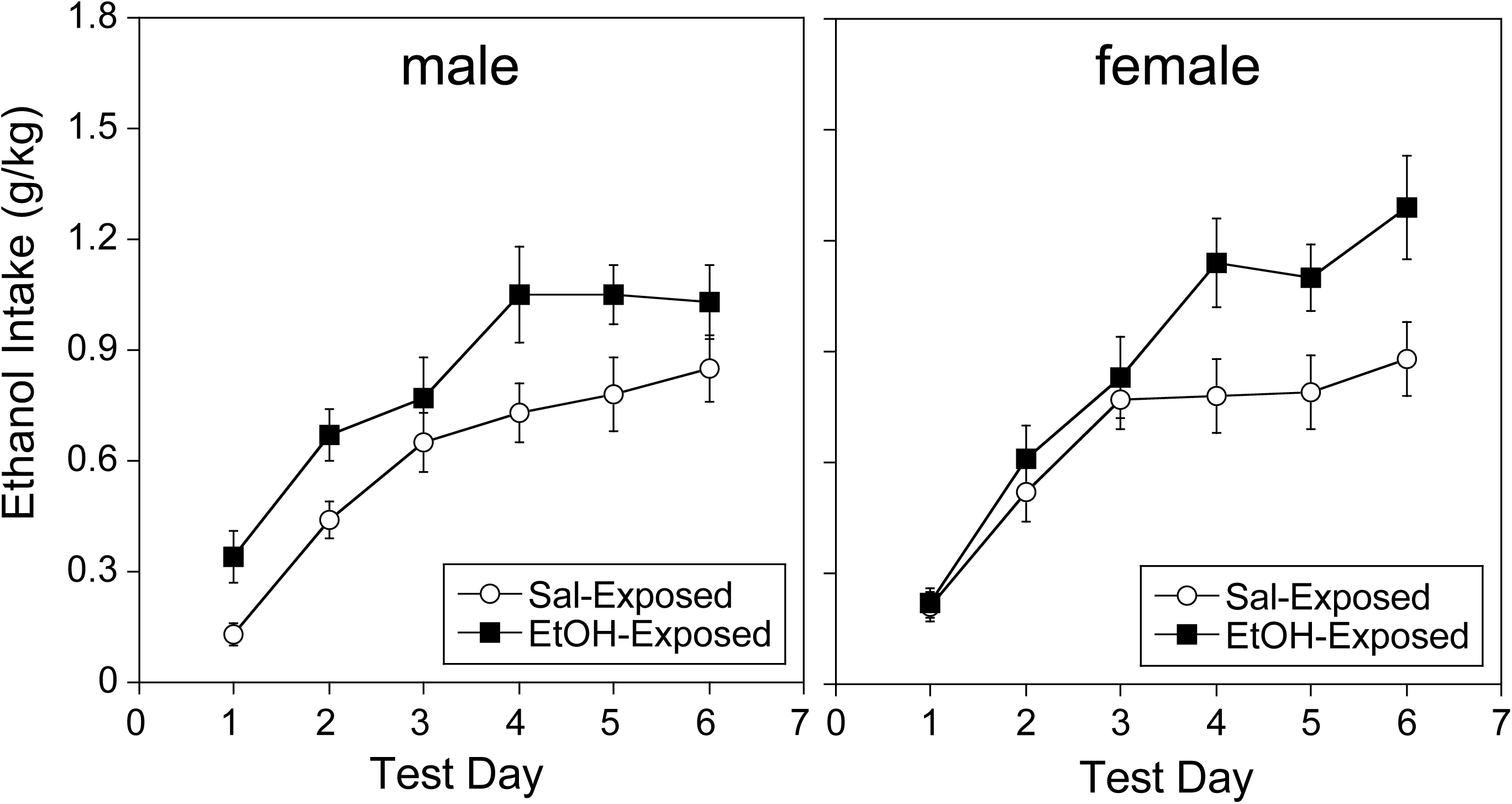
Ethanol intake in males and females: Impact of PAE. Prenatal exposure to ethanol significantly increases ethanol intake in adult males across all testing sessions (left). Conversely, prenatal ethanol exposure significantly increases ethanol intake in females in the last 3 sessions (right).

In males, BECs on the last drinking test day did not differ as a function of either exposure, F(1,36) = 0.022, p = 0.883, or context F(1,36) = 0.003, p = 0.958 (Figure 2, left). In contrast, ethanol intake regardless of context produced significantly higher BECs in ethanol-exposed females than in their saline-exposed counterparts, F(1,36) = 4,602, p < 0.05 (Figure 2, right).

**Figure 2:**
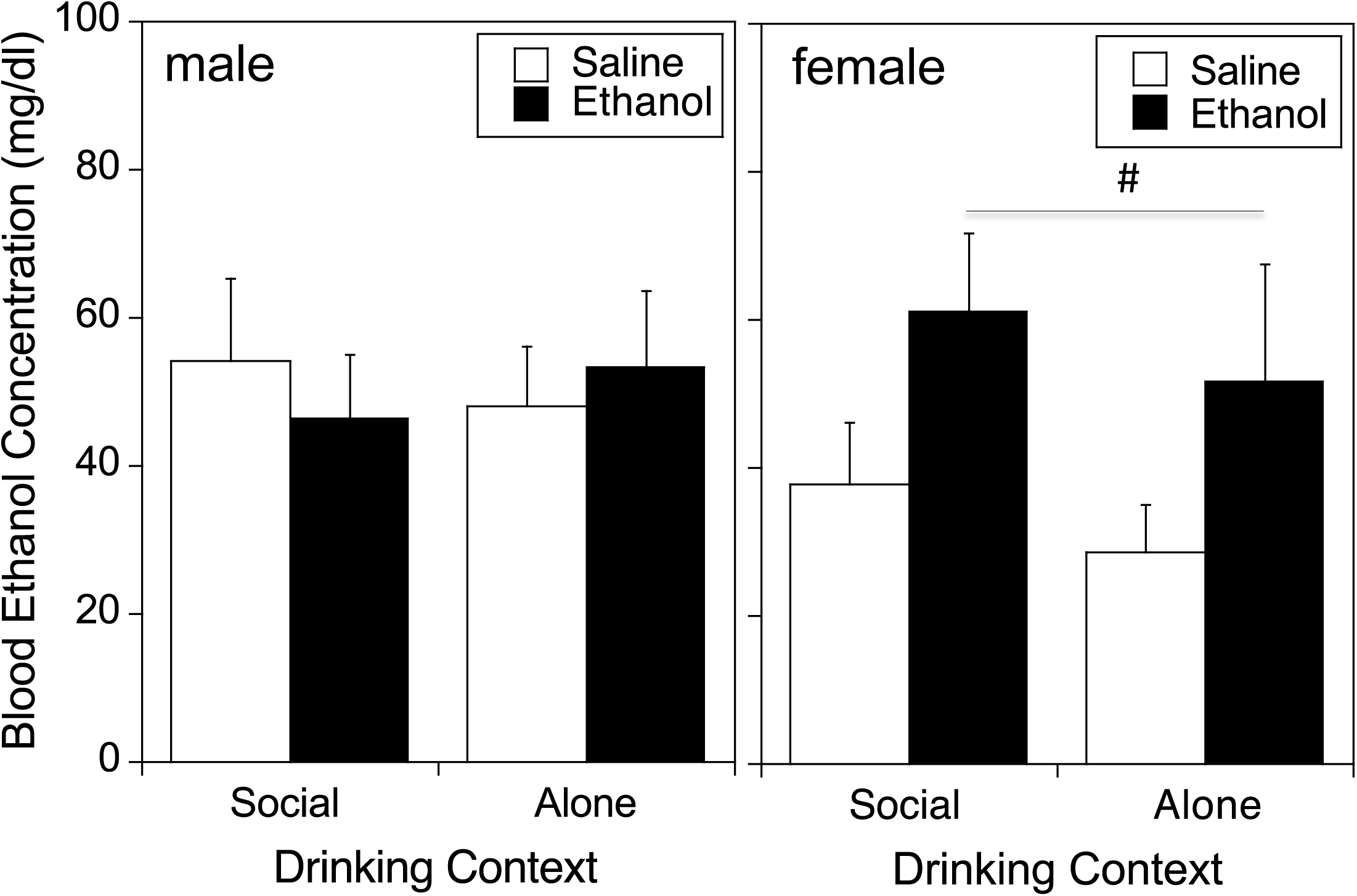
Blood ethanol concentrations (BECs) following final drinking session in males and females. Males (left) showed similar BECs across all groups, whereas PAE females (right) exhibited significantly (#, p <0.05) higher BECs relative to saline-exposed controls when collapsed across drinking context.

When the coefficient of social preference (an index of anxiety-like behavior) was used to divide males and females into tertiles and analyzed, in males, the coefficient differed only as a function of anxiety level, F(2,34) = 51.185, p < 0.0001, but not of exposure, F(1, 34) = 2.173, p = 0.149 (see Table 1), with ethanol- and saline-exposed males demonstrating similar social preference. In contrast, social preference in females was affected by exposure, F(1,34) = 17.603, p < 0.001, with ethanol-exposed females demonstrating significantly lower social preference than their saline-exposed control counterparts across all anxiety levels (see Table 1).

In saline-exposed control males, social anxiety level did not affect ethanol intake, F(2,17) = 0.484, p = 0.6243. However, saline-exposed males ingested more ethanol under social test circumstances than while drinking alone, as evidenced by a significant main effect of context, F(1, 17) = 8.282, p < 0.05 (see Figure 3). In ethanol-exposed males ethanol intake differed as a function of anxiety level, F(2,17) = 3.984, p < 0.05, with high socially anxious males ingesting more ethanol than their counterparts that demonstrated low and medium social anxiety levels (see Figure 3). However, there was no effect of context on ethanol intake in ethanol-exposed males. The analysis of ethanol intake in saline-exposed females revealed a significant anxiety level by context interaction, F(2,17) = 3.889, p < 0.05, with high socially anxious females drinking almost twice as much as their low and medium anxious counterparts under social drinking circumstances, while drastically decreasing their intake when drinking alone. Social anxiety levels did not affect ethanol intake of females prenatally exposed to ethanol under either drinking context.

**Figure 3:**
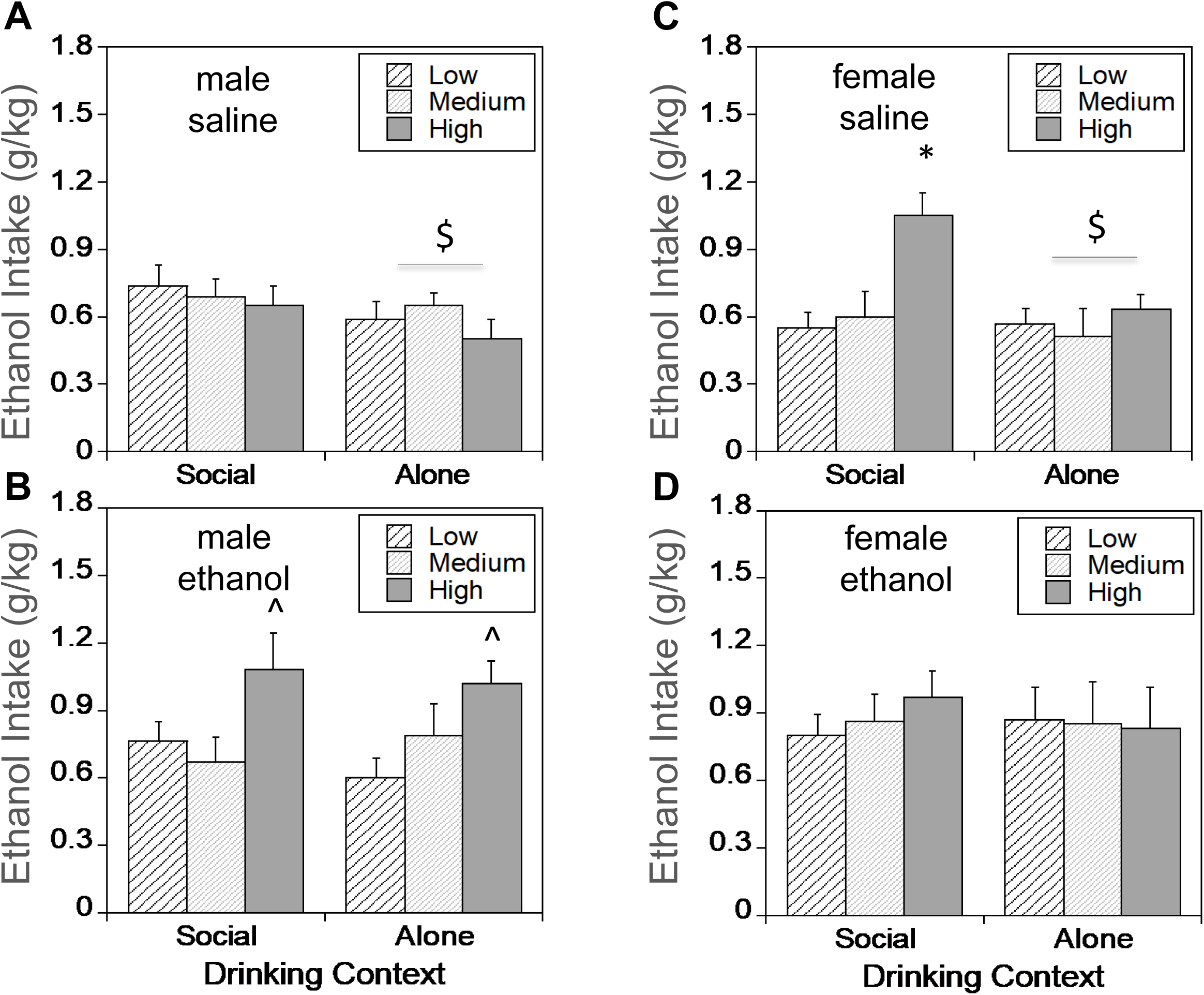
Ethanol intake under social or alone circumstances in males and females: Impact of PAE and levels of social anxiety-like behavior. (A) Ethanol intake was significantly lower in saline-exposed males when ethanol intake was tested alone relative to social drinking, regardless of social preference level ($ - p < 0.05 between drinking contexts). (B) In ethanol-exposed (PAE) males, ethanol intake was significantly higher in high socially anxious animals relative to low and medium socially anxious groups, with data collapsed across drinking context (^ - p < 0.05). (C) In control saline-exposed females, ethanol intake was significantly increased in high socially anxious animals compared to low and medium socially anxious groups under social drinking context (* - p < 0.05 compared to other groups within social drinking context; $ - p < 0.05 between drinking contexts). (D) Ethanol intake in ethanol-exposed (PAE) females did not differ among groups with different levels of social anxiety-like behavior or between drinking contexts.

## Discussion

Although PAE effects on social behavior and ethanol intake have been individually reported in separate studies, assessment of these behaviors has not been done in a within subject design. Consistent with previous reports, we found that a single exposure to high levels of ethanol on G12 increased ethanol intake and reduced social preference in adult female, but not male offspring, suggestive of heightened social anxiety-like behavior in these females. Surprisingly, our findings indicate that PAE male offspring that exhibited the lowest social preference and hence the highest levels of social anxiety consumed significantly more ethanol than their less anxious counterparts. Control males prenatally exposed to saline did not demonstrate a relationship between social preference and ethanol intake, while control females did, but only under social test circumstances. Additionally, whereas control male and female offspring consumed more ethanol in a social setting than alone, G12 PAE offspring of both sexes consumed the same amount of ethanol regardless of social context, which in general was more than saline-exposed offspring. Taken together, these data provide compelling evidence highlighting that PAE increases the relationship between social anxiety-like behavior in males and ethanol intake regardless of sex.

It is well-established that PAE significantly impacts social skills and interactions throughout the lifespan [53-57]. Alterations in social behavior have also been reported in numerous animal models [see review by [58]], although the direction of the effect is not always consistent. Several important factors that likely contribute to discrepant effects on social behavior include the timing and amount of ethanol exposure during gestation, in addition to the age of social testing and the testing assay. Furthermore, sex differences have not been consistently examined in neither clinical nor preclinical studies, providing another major factor in the interpretation of the literature. Nevertheless, we have identified G12 as a unique developmental epoch during which ethanol exposure can have long-lasting robust effects on social behavior [27, 44, 45, 59, 60]. PAE-induced social behavioral deficits are quite dynamic, with age- and sex-specific effects being apparent in the various components of social behavior, such as social investigation, social preference, contact, and play fighting. Importantly, social preference, which we have shown to reflect anxiety-like alterations evident under social circumstances, is consistently affected by high levels of G12 PAE in both sexes and at multiple age points. For example, in our last study using Sprague-Dawley rats, we found that social preference was reduced in juvenile and adolescent male offspring, but not in adult males, whereas females showed reduced social preference as adolescents and adults, but not as juveniles [27]. Consistent with our previous studies, in the current study we found that on average, social preference was unaffected in adult male PAE offspring, while adult females exhibited reduced social preference. However, when dividing subjects into level of social anxiety using a tertile split, we found that high levels of social anxiety contributed to high ethanol intake only in the PAE males. Although PAE females, in general, demonstrated increased ethanol intake and lower values of the coefficient (i.e., higher social anxiety) relative to saline-exposed controls, ethanol intake in these females did not differ as a function of social anxiety levels. In contrast, high socially anxious saline-exposed females ingested more ethanol under social circumstances than their less anxious counterparts and decreased ethanol intake while drinking alone.

PAE has been reported to increase the risk of substance use disorder, particularly regarding alcohol abuse. In contrast to adult lifetime prevalence rates of alcohol use disorders in the United States, estimated to be 10.1% in 2018 [61], it is estimated that ∼35% of individuals with an FASD have alcohol abuse problems [6-9, 62]. This PAE-induced predisposition for future alcohol abuse has been recapitulated in animal models, which has demonstrated that PAE across several gestational windows increases ethanol intake and preference at some point in the life-time of the offspring [28-35, 63]. Consistent with these findings, we found that that a single high level ethanol exposure on G12 increased ethanol intake in adult male and female offspring relative to control rats. It is worth noting that these differences were observed in adult offspring after only six drinking episodes, which is quite robust given that others have reported a lack of PAE effect on drinking in adults using a longer prenatal exposure (G17-20) and a much more extended ethanol intake paradigm (55 days) [63].

Although the results from the social behavior and ethanol intake were interesting and exciting given the brief level of PAE, the most novel and compelling finding from the current study was that the level of ethanol intake was highly associated with social anxiety-like behavior in PAE males. Specifically, our within-subject assessment revealed that PAE males with the highest level of social anxiety-like responses, although similar to those of saline-exposed controls, were high drinkers, whereas ethanol intake of low socially anxious PAE males was comparable to that of low socially anxious controls, suggesting that PAE-associated increases in ethanol intake are driven predominantly by high socially anxious individuals. As in our previous study[51], levels of social anxiety were not predictive of ethanol intake in saline-exposed control males. However, high socially anxious control females demonstrated the highest levels of social drinking comparable to those of PAE females, suggesting that these females might be more sensitive to anxiety-relieving effects of ethanol under social test circumstances. This is confirmed by a drastic decrease in ethanol intake demonstrated by these high socially anxious control females when they were tested alone.

Despite the clear association between social anxiety-like behavior and ethanol intake in PAE male offspring, we found that both male and female PAE offspring consumed the same level of ethanol when tested either alone or in a social environment (pair-housed). This was in contrast to control offspring that showed significantly higher ethanol intake in the social setting relative to drinking alone, an effect that was apparent in both males and females and consistent with what we have previously observed in ethanol naïve adults [51]. Importantly, these data indicate that despite social anxiety-like behavior contributing to high ethanol intake in PAE offspring, the drinking context (i.e. social or alone) does not influence the level of intake in PAE subjects.

While the behavioral alterations reported in this study are robust and novel, the underlying neurobiological mechanisms that contribute to these PAE effects are unknown. The neural networks known to drive social behavior are vast, but certain structures have been shown to regulate social behavior. For example, the importance of the amygdala in social processing is highlighted by studies examining the effects of amygdala damage in humans and nonhuman primates which reduces group formation, social judgement, trust acquisition, and interpretation of social cues [64-71]. Similarly, the medial prefrontal cortex has been shown to be highly involved in social processing and behavior [72-75]. More recently, it was shown that communication between the basolateral amygdala and the prefrontal cortex drives social interactions [76]. Importantly, there is evidence that both of these areas are altered by PAE [15, 18, 31, 33, 41, 77-80]. Furthermore, both the basolateral amygdala and the medial prefrontal cortex are highly implicated in alcohol abuse and alcohol use disorder [81]. Based on this, future studies should test the impact of G12 PAE on these neural networks to determine their contributions to the observed behavioral alterations reported in the current study.

These findings provide compelling evidence in support of anxiety-like (in this case social anxiety) driving ethanol intake and preference following PAE in males, a phenomenon which has been hypothesized and shown in prenatal ethanol-naïve individuals [10], but had not been demonstrated in PAE subjects. Furthermore, the subtle, but important sex-dependent social contributors to ethanol intake noted in this study shed light on our understanding of sex-dependent effects of prenatal alcohol, particularly related to long-term deficits in emotional processing and alcohol intake.

## Acknowledgements

This work was funded by NIAAA grant P50AA017823 and the Psychology department at Binghamton University.

